# An Integrated Physiological, Cytology and Proteomics Reveals Network of Sugarcane Protoplasts Responses to Enzymolysis

**DOI:** 10.1101/2022.09.30.510375

**Authors:** Demei Zhang, Rui Wang, Jiming xiao, Shuifang Zhu, Xinzhu Li, Shijian Han, Zhigang Li, Yang Zhao, Md. Jahidul Islam Shohag, Zhenli He, Suli Li

## Abstract

The protoplast experimental system has been becoming a powerful tool for functional genomics and cell fusion breeding. However, the physiology and molecular mechanism during enzymolysis is not completely understood and has become a major obstacle to protoplast regeneration. Our study used physiological, cytology, iTRAQ (Isobaric Tags for Relative and Absolute Quantification) -based proteomic and RT-PCR analyses to compare the young leaves of sugarcane (ROC22) and protoplasts of more than 90% viability. We found that oxidation product MDA content increased in the protoplasts after enzymolysis and several antioxidant enzymes such as POD, CAT, APX, and O^2-^ content significantly decreased. The cytology results showed that after enzymolysis, the cell membranes were perforated to different degrees, the nuclear activity was weakened, the nucleolus structure was not obvious, and the microtubules depolymerized and formed many short rod-like structures in protoplasts. The proteomic results showed that 1,477 differential proteins were down-regulated and 810 were up-regulated after enzymolysis of sugarcane young leaves. The GO terms, KEGG and KOG enrichment analysis revealed that differentially abundant proteins were mainly involved in bioenergetic metabolism, cellular processes, osmotic stress, and redox homeostasis of protoplasts, which would allow protein biosynthesis or / degradation. The RT-PCR analysis revealed the expression of osmotic stress resistance genes such as *DREB, WRKY, MAPK4*, and *NAC* were up-regulated. Meanwhile, the expression of key regeneration genes such as *CyclinD3, CyclinA, CyclinB, Cdc2, PSK, CESA* and *GAUT* were significantly down-regulated in the protoplasts. Hierarchical clustering, identification of redox proteins and oxidation products showed that these proteins were involved in dynamic networks in response to oxidative stress after enzymolysis. We used a variety of methods to figure out how young sugarcane leaves react to enzymes.

## 1. Introduction

Plant cells from which the cell wall has been enzymatically or mechanically removed are called protoplasts (Chupeau et al., 2013). Protoplasts can maintain the physiological and cellular processes of whole plants (X. Xu et al., 2021). Therefore, the protoplast experimental system is becoming a powerful tool for functional genomics in which to study protein–protein interactions, protein localization, and signaling pathways involved in plant physiology, innate immunity, growth and development (Page et al., 2019). In addition, somatic hybridization breeding from different species, therefore, supplies a practical breeding tool (Johnson & Veilleux, 2010) and circumvents sexual hybridization related prezygotic or postzygotic barriers. It can create different homokaryon or heterokaryon types, as well as alloplasmic hybrids (cybrids) (Xia, 2009). Somatic hybridization breeding has been reported for a wide variety of plant species, including rice (Ma et al., 2020), and maize (Zhou. B et al., 2008). However, regeneration is typically the bottleneck in somatic hybridization breeding programs, forcing researchers to develop more novel ways, such as the most appropriate donor material, source, optimal protoplast isolation, and appropriate culture method. (Davey et al., 2005).

Enzymolysis is the first step in somatic hybridization breeding, and the damage degree of protoplasts is the key link to determining the regeneration of protoplasts (Chupeau et al., 2013). To attain high regeneration rates, it was recommended that young and nonstressed tissues be used for protoplast isolation (Chupeau et al., 2013) and that plant growth conditions be adjusted to avoid premature cell death (Horii & Marubashi, 2005). However, plant protoplasts remain intact only at a proper osmotic pressure during enzymolysis and the osmotic pressure has to be adjusted in order to obtain the most viable protoplasts (Li. Z. G et al., 2020). The degradation of cell walls is performed in the presence of an osmotic, usually a sugar such as mannitol, sucrose, or sorbitol (Estavillo et al., 2014). This prevents protoplast disruption due to differences in osmolality between the cell interior and the digestion medium (Estavillo et al., 2014). The suggested link between osmolarity decrease or increase and chromatin over condensation may also contribute significantly to understanding the intricate interaction between genetic background and environmental factors in early regeneration (Chupeau et al., 2013).

When attempting the isolation of the cytosolic fraction from the plant, it is critical to start with a pure sample of intact protoplasts, as many break during digestion. Wounding resulting from the procedure to isolate protoplasts, and even the subsequent culture, has been found to be stressful to the cells and even suggested to be the trigger for dedifferentiation (T. Zhang et al., 2022). Oxidative stress evoked during protoplast isolation and their culture may be one of the factors contributing to the recalcitrance of protoplasts (Petřivalský et al., 2012). It has been reported that in rice, protoplasts can be damaged by reactive oxygen species (ROS) generated from cells treated with xylanase or pectin lyase (Ishii, 1988). The addition of superoxide dismutase and catalase to the rice protoplast isolation medium was found to improve protoplast viability (Pasternak et al., 2005). It is suggested that the different levels of reactive oxygen species (ROS) and antioxidant enzymes and scavengers present in citrus and cucumbers play a significant role in determining the regeneration capability of protoplasts derived from both types of cell (Petřivalský et al., 2012). Accumulated ROS are associated with increased levels of lipid peroxidation (Papadakis et al., 2001) and participate in the initiation of apoptosis-like cell death of cultured protoplasts (Yasuda et al., 2007).

The removal of the cell wall imposes a tremendous challenge to the cells. Consequently, plant cells respond to the removal of the cell wall in the nucleus at every level of the regulatory hierarchy (Mujahid et al., 2013). As a result, the enzymatic solution disrupts not only physiologic function but also protoplast cytology and molecule level (Fu, 2001). Isolation generates structural variability, resulting in anomalies observed during cell division (Tylicki, 2002), as well as in plants regenerated from protoplasts (Cambecedes et al., 1988). These anomalies often hinder the introduction of new plant varieties obtained by methods of in vitro protoplast fusion (Handley et al., 1986). Sections of protoplasts show that cortical microtubules are present at all times but examination of osmotically ruptured protoplasts (Marchant & Hines, n.d.). The early level opmental stages initiated in protoplasts are accompanied by large-scale chromatin remodeling (T. Zhang et al., 2022) and by major transcriptional changes (Hayat et al., 2022). Sharma et al. (2011) examined the transcriptome response to enzymatic removal of the cell wall (Sharma et al., 2011). They found that kinases, transcription factors and genes predicted to be involved in cell wall-related functions were enriched in the differentially regulated gene category (Mujahid et al., 2013). The nuclear proteome differential expression in response to removal of the cell wall in rice suspension cells, including transcription factors, histones, histone domain containing proteins, and histone modification enzymes (Mujahid et al., 2013). Gene ontology analysis of the differentially expressed proteins indicates that chromatin & nucleosome assembly, protein-DNA complex assembly, and DNA packaging are tightly associated with cell wall removal (Mujahid et al., 2013). The osmotic stress effect at the time of protoplast separation altered the expression of some of the resistance genes, leading to browning (Ondřej et al., 2009). Little is known about the mechanisms that plant cells use to sense enzymatic removal of the cell wall and transduce corresponding signals to regulate cellular responses to maintain protoplast integrity. A greater application of molecular biology techniques can increase our understanding of plant cryobiology. By combining physiology, cytology and molecular biology approaches, an array of methods are available to understand how cells are protected during the enzymolysis process.

Sugarcane (*Saccharum spp.*) is an industrially major crop in tropical and sub-tropical areas, which contributes 70% global sugar production. It is also utilized for biofuel, ethanol and other by-products formation such as paper, plywood, animal feed, and industrial enzymes (Sheen, 2001). Somatic hybridization in sugarcane enables us to broaden the germplasm base. The current research on sugarcane protoplasts has mainly focused on optimizing enzymolysis conditions such as enzymolysis method, medium type, hormone type and concentration for protoplasts. The highly viable sugarcane protoplasts obtained by enzymolysis of young sugarcane leaves using the optimal mannitol concentration showed severe browning at a later stage and the cells were unable to divide continuously, which greatly hindered the regeneration of sugarcane protoplasts (Meena et al., 2022). If the problem of late protoplast browning is not solved, it will make the regeneration of sugarcane protoplasts a major problem (Meena et al., 2022). In fact, sugarcane protoplasts are inevitably affected by external conditions during enzymolysis, thus affecting the expression of relevant genes and their regeneration ability (Tessadori et al., 2007). In our previous report, transcriptome sequencing was performed on young sugarcane leaves and protoplasts after enzymolysis, and we found significant difference in uniqueness, differentially expressed genes and the DEGs (Differentially expressed genes) categories (D. Zhang, et al., 2022). However, the molecular mechanism of how protein regulatory networks regulate sugarcane protoplast cell division, differentiation, and further rooting into seedlings remains unknown. Integrated cytology, physiology and proteomics analysis of the response of sugarcane young leaves to enzymolysis will provide key information as to the timing and nature of the recovery pathways in systems that survive or fail to enzymolysis. In this study, we used physiology, cytology, proteomics, and RT-PCR ((Reverse Transcription-PCR)) to examine differences in sugarcane protoplasts before and after enzymolysis, functional enrichment analysis of differential genes, and functional annotation of protein to provide an answer to the molecular, physiological, and cytology mechanisms underlying the difficulties in regenerating sugarcane protoplasts. This study provides criteria and a theoretical foundation for the construction of a standard system for producing regenerated protoplasts through the use of molecular markers and antibody detection of enzymolysis.

## 2. Result

### 2.1 Effect of enzymolysis on subcellular structure of sugarcane protoplasts

The young sugarcane leaf cells (Fig. 1a) were rounded after enzymatic digestion and had smooth cell membranes (Fig. 1b, f). The Fluorescein diacetate (FDA) detection results showed that the viability of the enzymatically digested sugarcane protoplasts activity was above 90% (p < 0.05) (Figure 1c.). The cell membranes of young sugarcane leaves were intact before enzymolysis (Fig. 1d), however, the cell membranes of protoplasts perforated to different degrees after enzymolysis (Fig. 1e).. The results of DAPI fluorescence staining showed that the nucleolus of nuclear membrane was clear and DAPI fluorescence was strong before enzymolysis (Fig. 1g,h). The nuclear membrane of protoplasts was still intact after enzymatic hydrolysis, but the blue fluorescence and nuclear activity were weakened and the nucleolus structure was not obvious (Fig. 1i). There was a tight association between microtubules and the plasma membrane in young sugarcane cells, the microtubule array was well organized (Fig. 1 j,k.), However microtubules depolymerize and formed many short rod-like structures in protoplasts, and a large number of adhesions of periplasmic microtubules were observed on the plasma membrane of newly isolated protoplasts in a radial or fan-shaped distribution (Fig. 1 l.).

**Figure 1.**
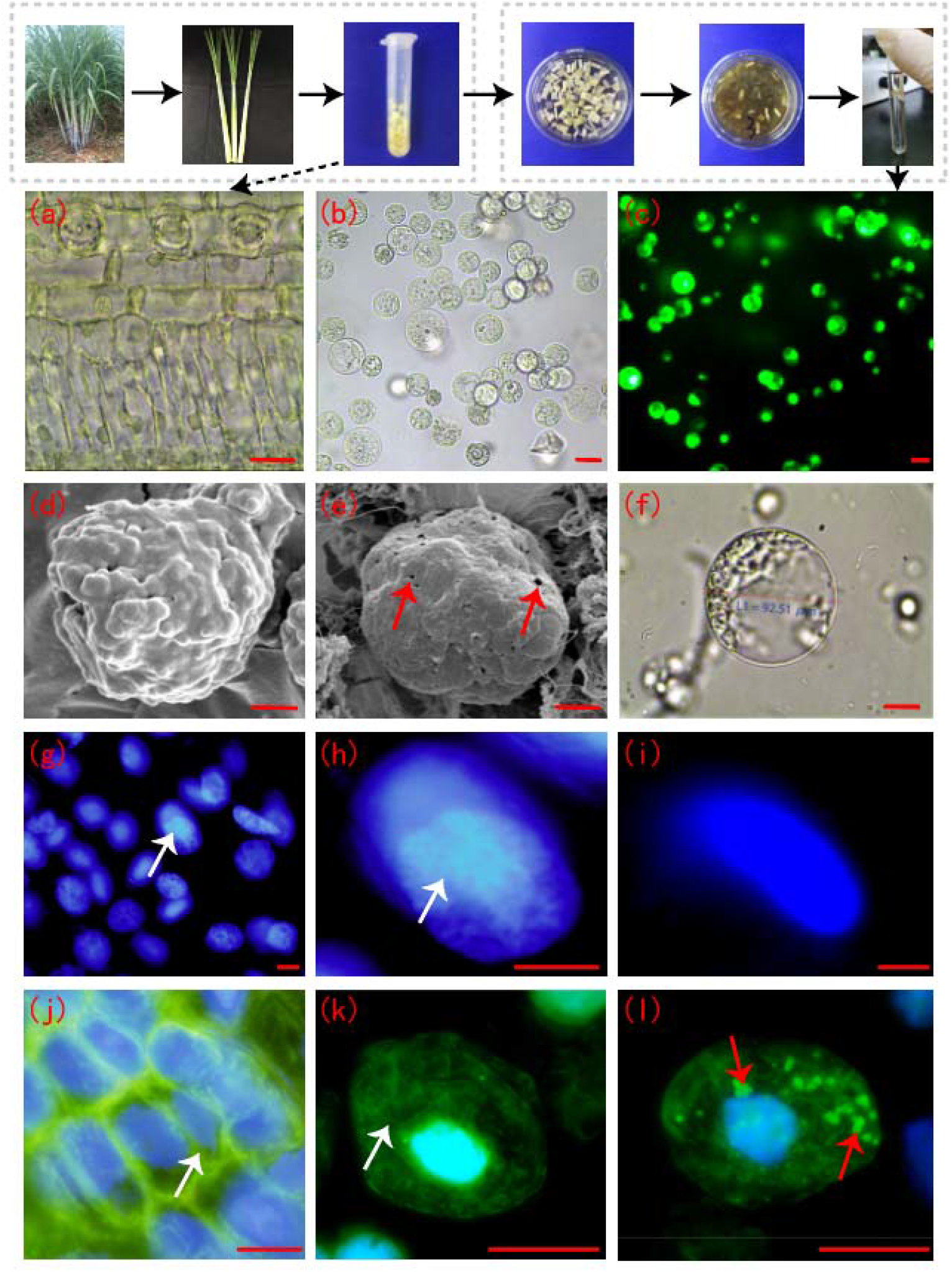
Sugarcane protoplasts after enzymolysis. (**a**) Cellular morphology of young sugarcane leaves before enzymolysis, the cells were arranged neatly and structure was intact, bar = 30μm; (b) Sugarcane protoplasts obtained after enzymolysis, Protoplasts were round, full and smooth, bar = 25μm; (**c**) FDA detected the vitality of protoplasts, FDA had strong fluorescence and high cell viability, bar = 25μm; (d) Cell membrane of young sugarcane leaves under Scanning Electron Microscope, cell wall integrity, bar = 3μm; (e) Cell membrane of sugarcane protoplasts under Scanning Electron Microscope, there were little perforations in the membrane, Arrows indicate perforations, bar = 3μm; (f) Cell membrane morphology of sugarcane protoplasts under Light Microscope, expanded protoplasts with smooth and intact cell membranes, bar = 20μm; (**g, h**) Nuclei of sugarcane young leaf cells after DAPI,the nucleolus and nuclear envelope are clear, the arrows indicate nucleolus,, bar = 5μm; (i) Nuclei of protoplasts after DAPI, the nucleolus ware not clear bar = 5μm; (j, k) Periplasmic microtubule array in young sugarcane leaf cells with neat and orderly microtubule arrangement, the arrows indicating microtubule array, bar = 3μm; (l) Microtubule array of protoplasts, form many short rod-like structures, the arrows indicating microtubule depolymerization, bar = 3μm. **Note: (j-l) Green is FITC fluorescence, showing microtubulin; blue fluorescence is DAPI fluorescence, showing the nucleus.**

### 2.2 Effect of protoplasts by oxidative stress during enzymolysis

After enzymolysis, the content of MDA (Malondialdehyde) in protoplasts was significantly increased, and it was 4.7 times higher than that of young leaves (Fig. 2a), while the content of O^2-^ in protoplasts was significantly lower and only 1.2% of young leaves (Fig. 2b). There was no significant difference in SOD (Superoxide Dismutase) activity between sugarcane young leaves and protoplasts (Fig. 2c). However, compared with young leaves of sugarcane, contens of POD (Peroxidase), CAT (Catalase), and APX (Ascorbic Acid Peroxidase) in protoplasts were significantly reduced to 17.7%, 6.5%, and 17.5%, respectively (Fig. 2d, e, and f.).

**Figure 2.**
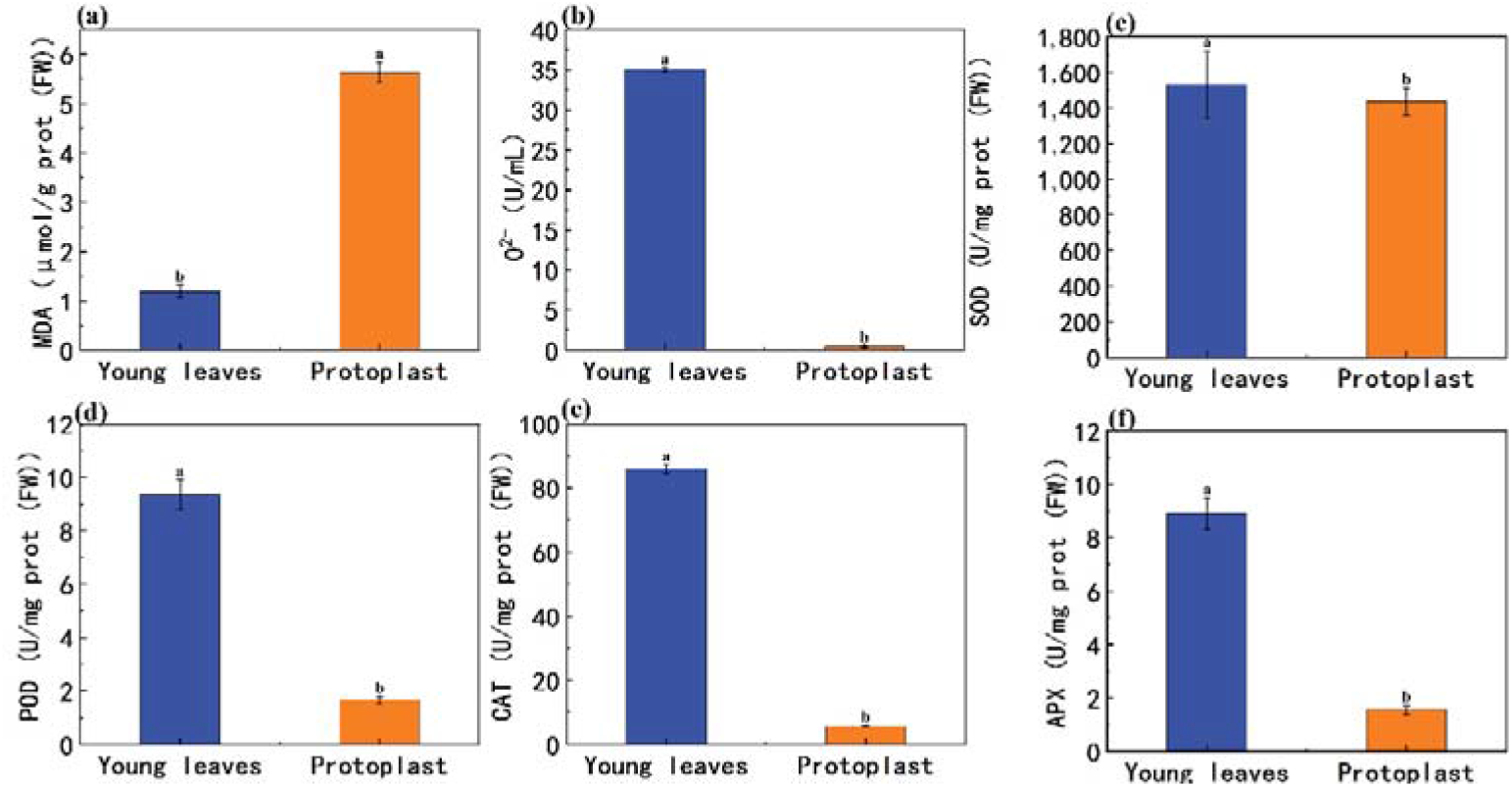
Expression of on oxidation products and antioxidant enzymes between sugarcane young leaves and protoplasts. (a) After enzymolysis, MDA content increased to 4.7 times that of young leaves; (b) After enzymolysis, the content of O^2-^ in protoplasts was only 1.2% of that in young leaves; (c) Compared with young sugarcane leaves, SOD content decreased to 93.7% after enzymolysis; (d) Compared with young sugarcane leaves, POD content decreased to 17.7% after enzymolysis; (e) Compared with young sugarcane leaves, CAT content decreased to 6.5% after enzymolysis; (f) Compared with young sugarcane leaves, APX content decreased to 17.5% after enzymolysis.

After enzymolysis, Gu/ZnSOD and CAT expression levels in protoplasts were down-regulated compared with young leaves, which accounted for only 1.6% and 2.8% of that of young leaves, respectively(Fig. 3).

**Figure 3.**
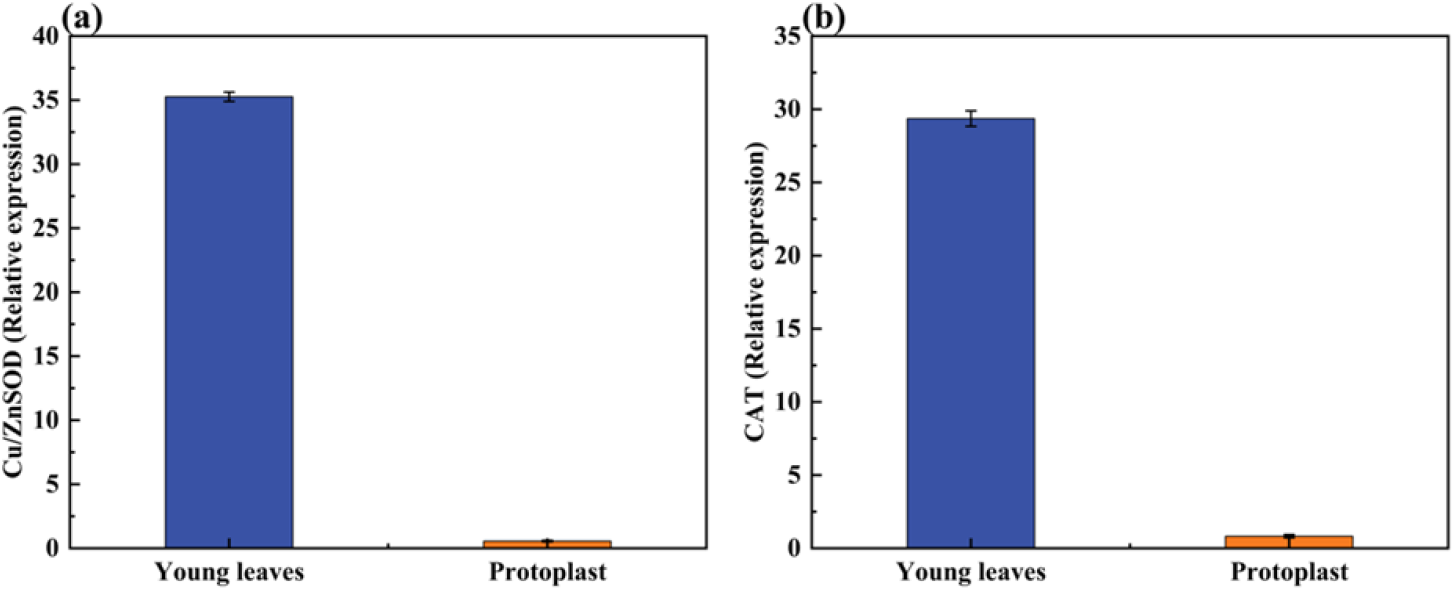
Expression of oxidase-related genes after enzymolysis. (a) After enzymolysis, the expression of Gu/ZnSOD was only 1.6% of that in young leaves; (b) After enzymolysis, the expression of CAT was only 2.8% of that in young leaves.

### 2.3 Effect of enzymolysis on expression of genes associated with osmotic stress

The expression of *DREB, WRKY, MAPK4* and *NAC* genes in sugarcane young leaves and protoplasts after enzymolysis were detected by fluorescence quantitative PCR with *GADPH* as internal reference genes. As shown in the figure, the expression of DREB, WRKY, MAPK4 and NAC genes in protoplasts were significantly up-regulated, and were 21 fold, 57,184 fold, 6,100 fold, and 200,050 fold higher than that of sugarcane young leaves, respectively (Figure 4a-d).

**Figure 4.**
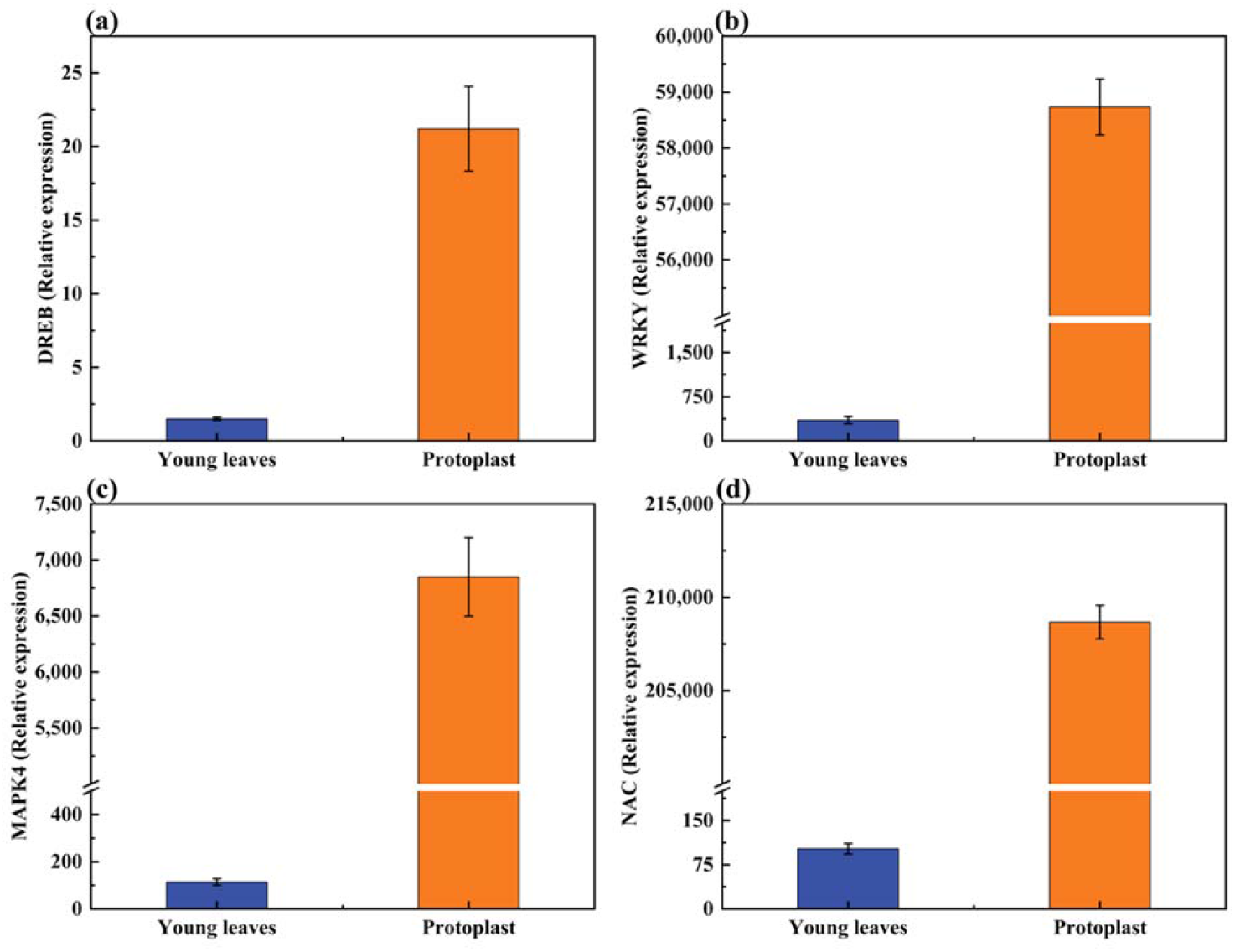
Expression of genes associated with osmotic stress after enzymolysis. Let the expression of young leaves be 1. **(a)** After enzymolysis, *DREB* expression quantity increased to 21 times that of young leaves; (b) After enzymolysis, *WRKY* expression quantity increased to 57,184 times that of young leaves; (c) After enzymolysis, *MAPK4* expression quantity increased to 6,100 times that of young leaves; (d) After enzymolysis, *NAC* expression quantity increased to 200,050 times that of young leaves.

### 2.4 Overview of proteome between sugarcane young leaves and protoplasts

Proteins with a fold change>=2 and pvalue<0.05 were significantly differentially expressed proteins (DEPs). The 2,287 DEPs in sugarcane protoplasts were identified through statistical analysis of young leaves and protoplasts after enzymatic digestion. Of them, 810 were up-regulated while 1,477 were down-regulated (Fig. 5A). In addition, Figure 6. showed that six samples were large and the experimental data were effective and reasonable.

**Figure 5.**
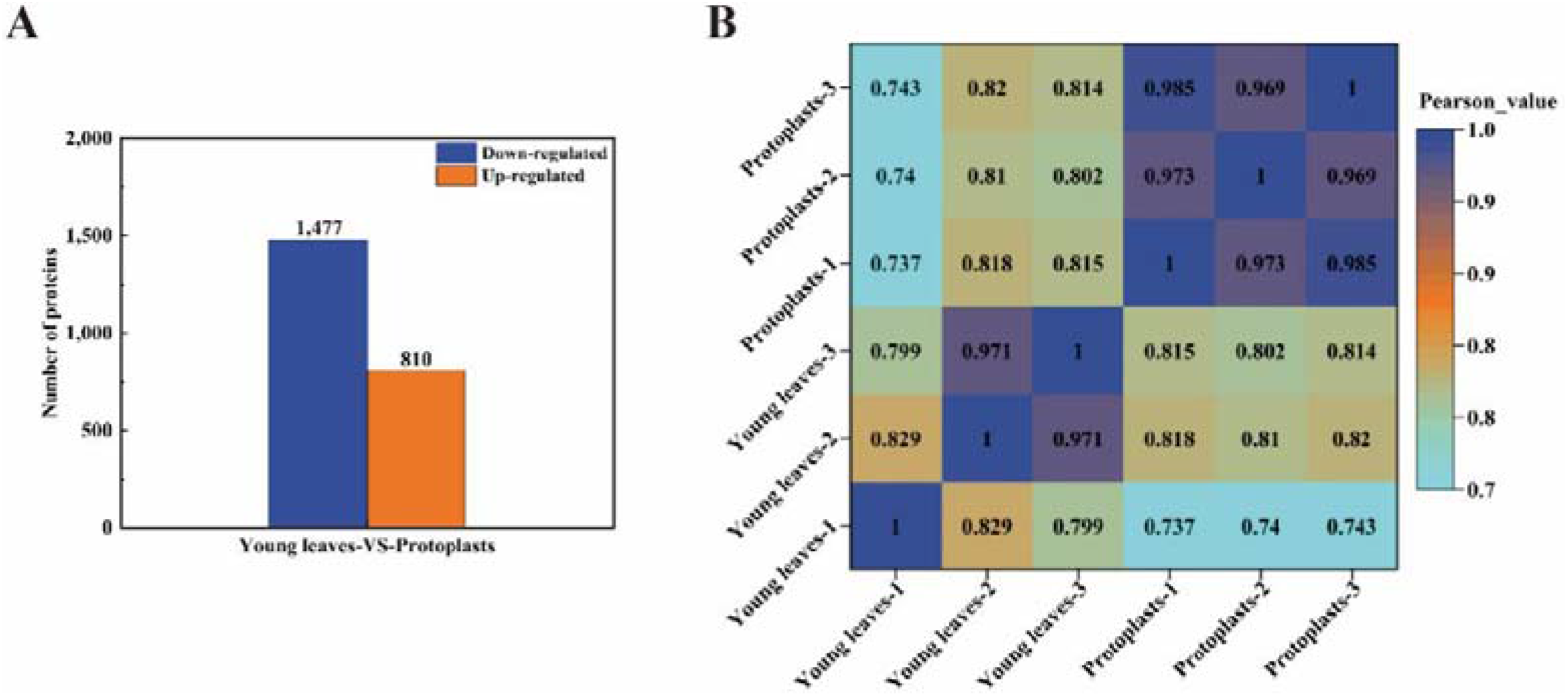
(A): Differentially expressed proteins between sugarcane young leaves and protoplasts; (B) Heat map of sample correlation analysis.

**Figure 6.**
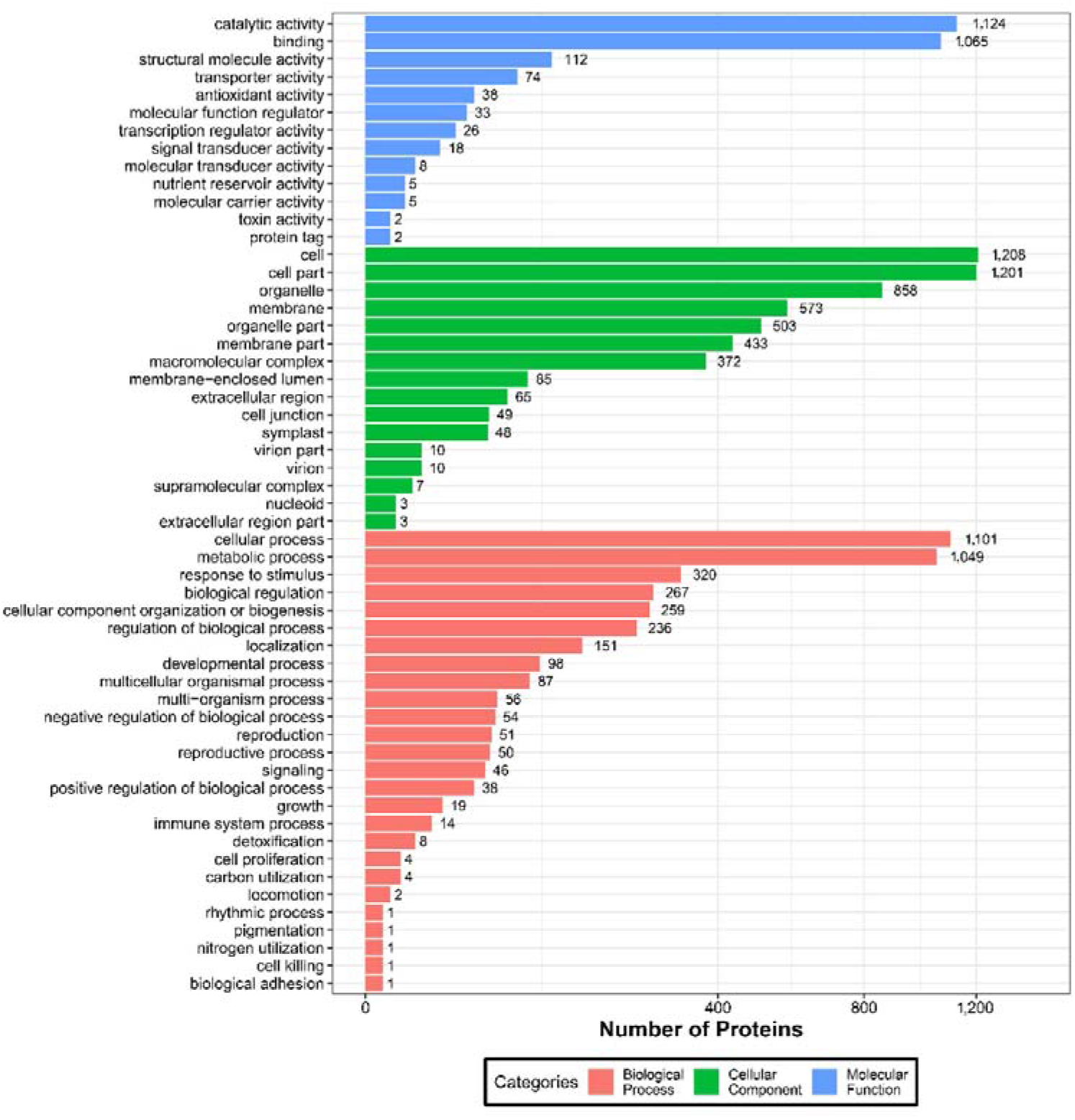
GO analysis of DEPs with young leaves VS protoplasts

Wolf PSORT software was used to locate the subcellular differential protein between sugarcane young leaves and protoplasts, and subcellular location of all identified DEPs showed that 839 proteins (36.9%) were located in the chloroplasts, 574 (25.3%) in the cytochylema, 444 (19.5%) in the nucleus, 164 (7.2%) in the Plasmalemma, 103 (4.6%) in the Mitochondria, 46 (2.0%) extracellular, 32 (1.4%) in the Tonoplast, 31 (1.4%) in the ER, 28 (1.2%) in the Cytoskeleton, 12 (0.5%) in the Peroxisome. Thus, many proteins including energy metabolism are greatly affected by enzymolysis (Table 1).

**Table 1.**
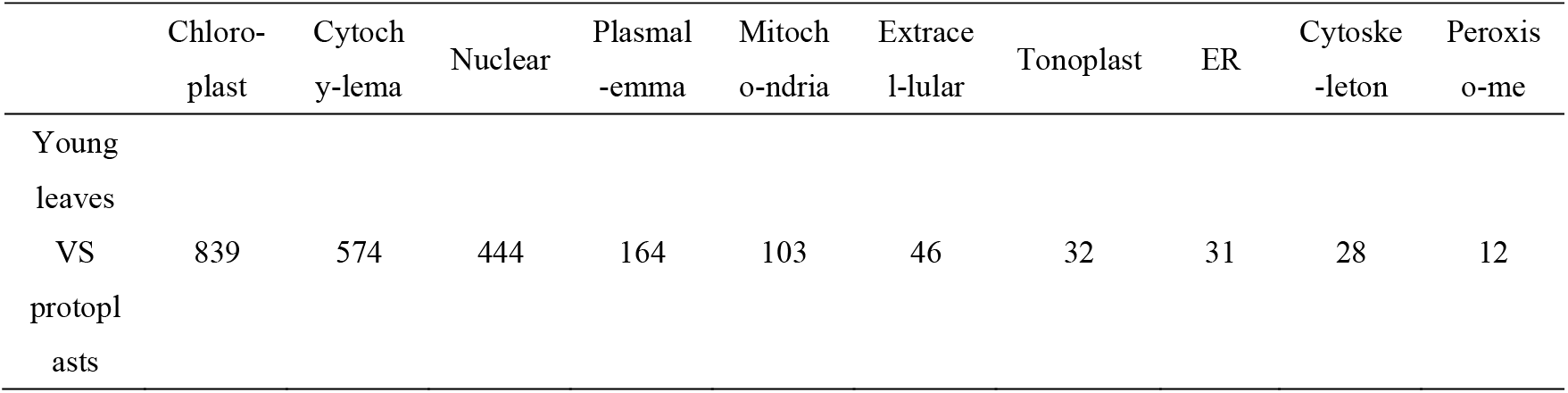
Subcellular localization of DEPs.

#### 2.5.1 Gene ontology (GO) classification of DEPs

To further understand the identified and quantified DEPs, we annotated their functions and features using GO enrichment analysis. The DEPs were grouped into three hierarchically structured GO terms: biological process, molecular function and cellular component (Fig. 6.). There were significant differences in GO terms between sugarcane young leaves and protoplasts. The DEPs were highly enriched in catalytic activity, cellular and metabolic process. The molecular function ontology analysis was as follows: catalytic activity (1,124 proteins), bingding (1,065 proteins), structural molecule activity (112 proteins), transporter activity (74 proteins) antioxidant activity (38 proteins) and so on. In the cellular component, the protein distributions were enriched in the cell (1,208), cell part (1,201 proteins), organelle (858 proteins), membrane (537 proteins), organelle part (503 proteins) and membrane part (433 proteins). The main biological processes of the differential expression proteins seem to be in cellular process (1,101 proteins), metabolic process (1,049 proteins), response to stimulus (320 proteins), biological regulation (267 proteins), cellular component organization or biogenesis (259 proteins) and so on. According to biological process classification, most of the proteins are related to cellular process and metabolic process. These DEPs were mainly located in the chloroplasts, cytochylema, nucleus, plasmalemma, and mitochondria. As can be seen from Figure 2, most of the DEPs in these pathways are in a down-regulated state, which indicates that they negatively regulates sugarcane protoplasts during enzymolysis.

#### 2.5.2 Kyoto encyclopedia of genes and enomes (KEGG) pathway analysis of DEPs

We used the KEGG database to identify enriched pathways using a two-tailed Fisher’s exact test to determine the enrichment of DEPs against all identified proteins (p-value < 0.05). KEGG cluster analysis between sugarcane young leaves and protoplasts showed that proteins were enriched in 7 pathways (Table 3–5), including 604 metabolic pathways (29.19%), 120 carbon metabolism (5.8%), 119 ribosome (5.75%), 110 biosynthesis of amino acids (5.32%), 80 protein processing in endoplasmic reticulum (3.78%), 77 glycolysis / gluconeogenesis (3.72%) and 72 spliceosome (3.48%). Except for glycan biosynthesis and metabolism and lipid metabolism, the other significant enrichment pathways were down-regulated more than up-regulated after enzymolysis. It can be seen that enzymolysis has a negative regulation on sugarcane protoplasts. Total significant enrichment KEGG pathways were listed in Table 2.

**Table 2.** Significant enrichment pathway of KEGG with Young leaves VS protoplasts.

| The main classification | Pathway | Diff Proteins with pathway annotation (2,069) | All Proteins with pathway annotation (7,650) | Up | Down |
| --- | --- | --- | --- | --- | --- |
| Amino acid metabolism | Arginine and proline metabolism | 24 (1.16%) | 47 (0.61%) | 10 | 14 |
|  | Cysteine and methionine metabolism | 40 (1.93%) | 109 (1.42%) | 6 | 34 |
| Carbohydrate metabolism | Glycolysis / Gluconeogenesis | 77 (3.72%) | 175 (2.29%) | 20 | 57 |
|  | Fructose and mannose metabolism | 35 (1.69%) | 76 (0.99%) | 9 | 26 |
|  | Pentose phosphate pathway | 31 (1.5%) | 70 (0.92%) | 9 | 22 |
|  | Ascorbate and aldarate metabolism | 22 (1.06%) | 51 (0.67%) | 7 | 15 |
|  | Galactose metabolism | 28 (1.35%) | 69 (0.9%) | 8 | 20 |
| Protein synthesis | Ribosome | 119 (5.75%) | 218 (2.85%) | 31 | 86 |
|  | Protein processing in endoplasmic reticulum | 80 (3.87%) | 190 (2.48%) | 43 | 37 |
|  | Spliceosome | 72 (3.48%) | 198 (2.59%) | 10 | 62 |
| Energy metabolism | Photosynthesis - antenna proteins | 9 (0.43%) | 9 (0.12%) | 9 | 0 |
|  | Carbon fixation in photosynthetic organisms | 42 (2.03%) | 96 (1.25%) | 18 | 24 |
|  | Photosynthesis | 20 (0.97%) | 37 (0.48%) | 19 | 1 |
|  | Nitrogen metabolism | 15 (0.72%) | 33 (0.43%) | 4 | 11 |
| Metabolism of<br>other amino acids | Glutathione metabolism | 41 (1.98%) | 100 (1.31%) | 10 | 31 |
| Global and<br>overview maps | Carbon metabolism | 120 (5.8%) | 315 (4.12%) | 39 | 81 |
|  | Biosynthesis of amino acids | 110 (5.32%) | 295 (3.86%) | 18 | 92 |
|  | Metabolic pathways | 604 (29.19%) | 2016 (26.35%) | 234 | 368 |
| Glycan<br>biosynthesis and<br>metabolism | N-Glycan biosynthesis | 21 (1.01%) | 41 (0.54%) | 20 | 1 |
| Lipid metabolism | Fatty acid elongation | 13 (0.63%) | 28 (0.37%) | 12 | 1 |
|  | alpha-Linolenic acid<br>metabolism | 24 (1.16%) | 62 (0.81%) | 19 | 5 |

#### 2.5.3 KOG analysis of DEPs

The potential function of DEPs of young leaves and protoplast after enzymolysis was analyzed comprehensively by KOG, and the results of KOG analysis were listed (Fig. 7). Among these DEPs, 337 were general function prediction only proteins and 99 were functionally unknown. The others were located in carbohydrate transport and metabolism (167), energy production and conversion (120), amino acid transport and metabolism (119), secondary metabolism biosynthesis, translation, ribosomal structure and biogenesis (215), RNA processing and modification (129), posttranslational modification, protein turnover, chaperones (273), signal transduction mechanisms (154), intracellular trafficking, secretion, and many in cell wall/ membrane/ envelope biogenesis, cell cycle control, cell division, chromosome partitioning, cytoskeleton and so on, which were of great significance for resolving the mechanism of the protoplast.

**Figure 7.**
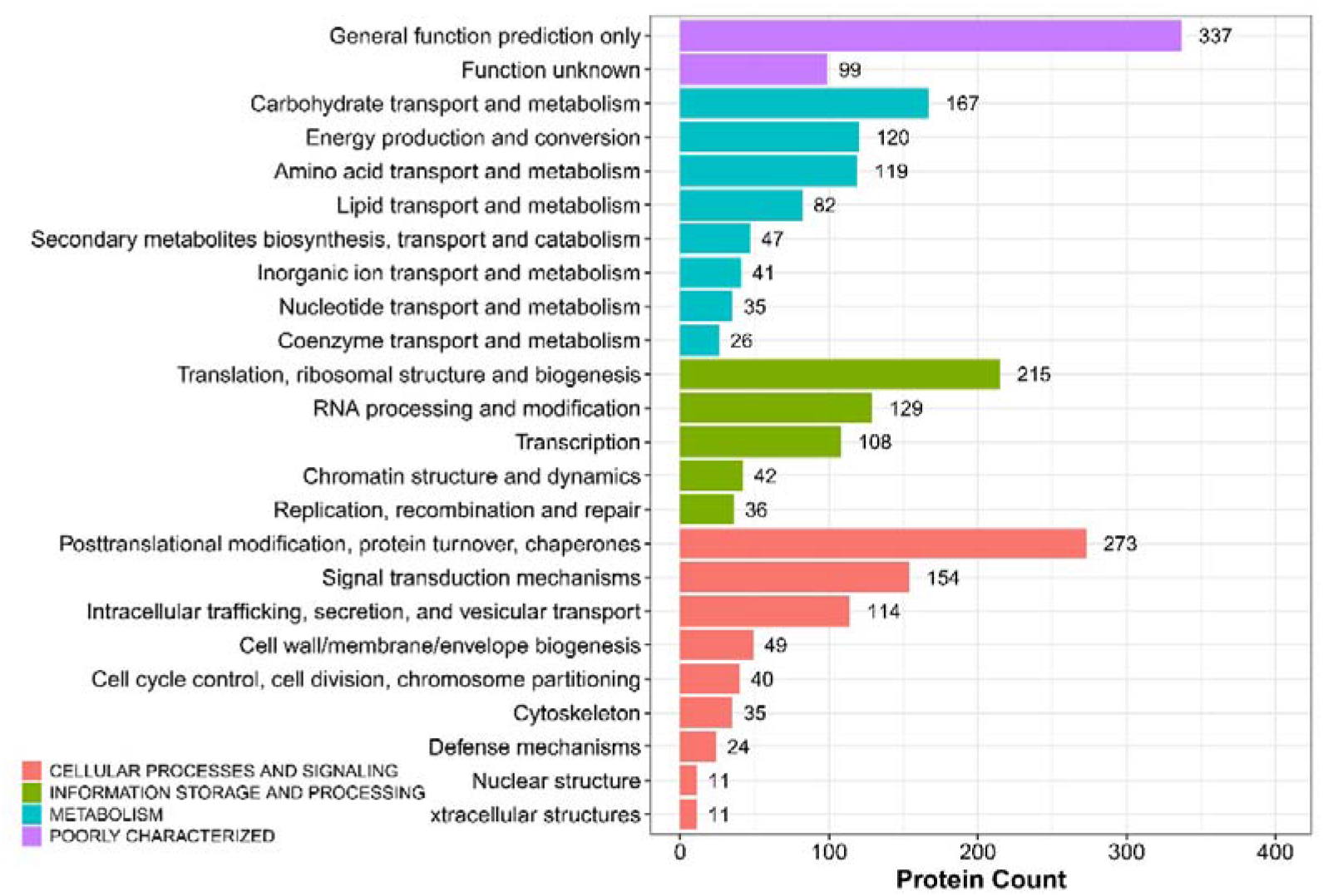
KOG analysis of DEPs with young leaves VS protoplasts

#### 2.5.4 Differential abundance of energy metabolism proteins and cellular process protei ns

We identified 54 candidate DEPs associated with energy metabolism in young leaves VS protoplastsafter enzymolysis. Of them, 22 were up-regulated while 32 were down-regul ated (Supplementary Table S1).

A total of 12 candidate DEPs associated with cell walls were obtained in young leaves and protoplasts. Of them, six were down-regulated, mainly including chitinase and 4,6-dehydratase/UDP-glucuronic acid decarboxylase. Acetylglucosaminyl transferase EXT1, apolipoprotein D/Lipocalin and pectin acetylesterase and similar were main up-regulated candidate DEPs (Supplementary Table S2).

A total of 12 candidate DEPs associated with cell cycle were obtained in young leaves VS protoplast after enzymolysis. Seven DEPs were down-regulated, including zw10 (tr|A0A194YN98|A0A194YN98_SORBI), late promoting complex, Cdc20, Cdh1, Ama1 subunits (tr|A0A1D6IIL4|A0A1D6IIL4_MAIZE), apoptosis-associated proteins/predictive DNA-binding proteins (tr|A0A1D6FJH1|A0A1D6FJH1_MAIZE), Microtubule-associated proteins essential for late spindle elongation MAP65-1a (tr|A0A1D6LRY4|A0A1D6LRY4_MAIZE), cell cycle associated protein Mob1-1 (tr|A0A059PYU0|A0A059PYU0_9POAL), ATM/Tel1(tr|A0A096SC75|A0A096SC75_MAIZE). While proteins associated with anti-cell death (tr|A0A096SC75|A0A096SC75_MAIZE) and proteins predicted to be involved in the formation of spindle matrix (tr|C5X7T2|C5X7T2_SORBI) were up-regulated (Supplementary Table S3).

#### 2.5.5 Differentially Abundant Secondary Metabolites Proteins

We identified seven candidate DEPs associated with the synthesis of secondary metabolites young leaves and protoplasts, which were mainly classified into biosynthesis of scop olamine, pethidine and pyridine alkaloids and styrene acrylic. Only protein ECERIFERUM 26-like was up-regulated among them, while Aspartate aminotransferase/Glutamic oxaloaceti c transaminase (AAT1/GOT2), Cytochrome P450 CYP2 subfamily, Alcohol dehydrogenase, Agmatine coumaroyl transferase-2, and Flavonol reductase/cinnamoyl-CoA reductase (Supple mentary Table S4).

#### 2.5.6 Differentially Abundant Antioxidant Proteins

In sugarcane young leaves and protoplast, 54 candidate DEPs linked with antioxidants were identified. Most of them were down-regulated, including ascorbate peroxidase, glutathione peroxidase, peroxidase, catalase, NADP-dependent isocitrate dehydrogenase, glutathione S-transferase and so on. While 3-oxoacyl CoA thiolase, glutaryl-CoA dehydrogenase, long-chain acyl-CoA synthetases (AMP-forming) and peroxisomal membrane protein MPV17 and related proteins were up-regulated (Supplementary Table S5).

### 2.6 Effect of enzymolysis on expression of regeneration -related genes after enzymolysis

**Figure 8.**
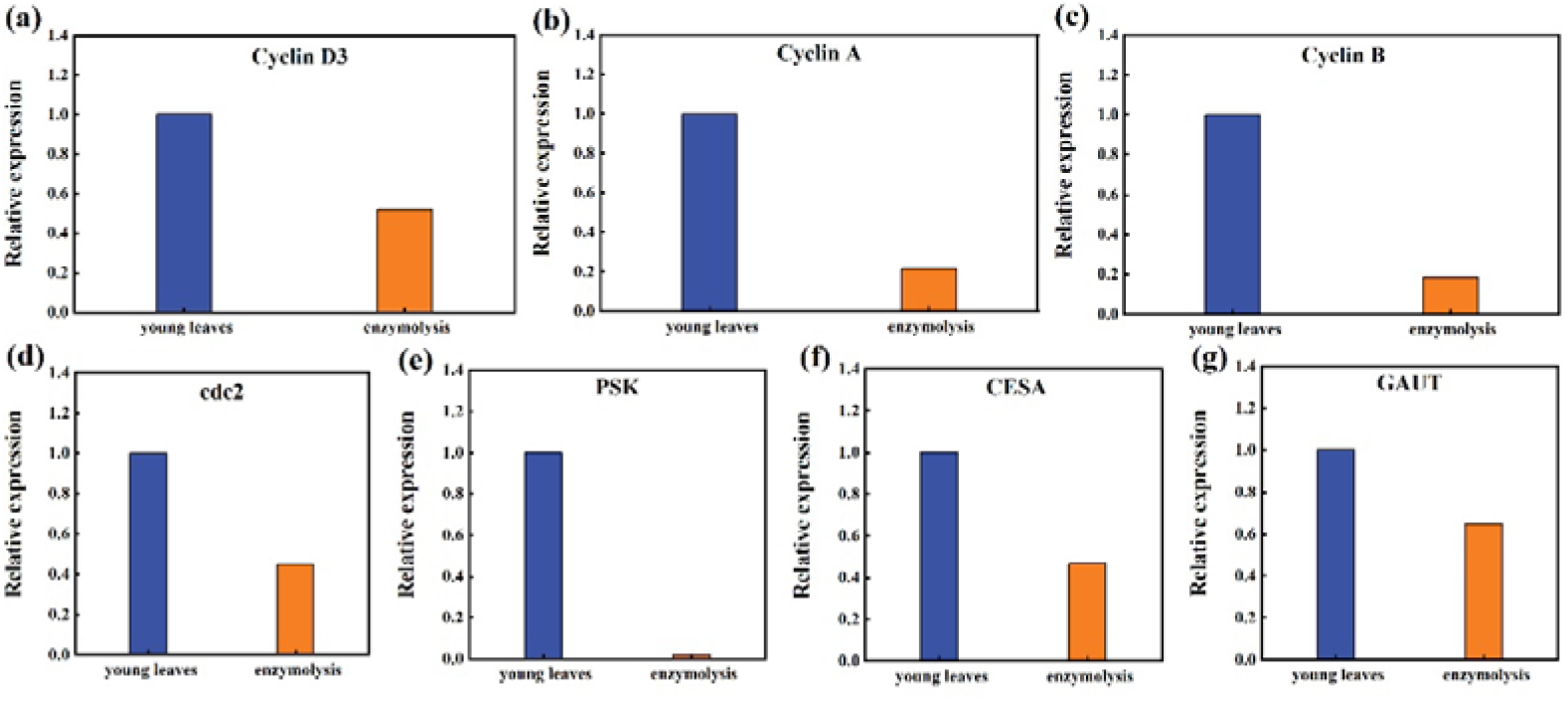
Expression of *Cyclin D3, Cyclin A, Cycliin B, cdc2, PSK, CESA, GAUT* genes between young leaves and protoplasts. (a) The expression of *Cyclin D3* after enzymolysis was only 52% of that of young leaves; (b) The expression of *CyclinA* after enzymolysis was only 21.32% of that of young leaves; (c) The expression of *CyclinB* after enzymolysis was only 18.60% of that of young leaves; (d) The expression of *cdc2* after enzymolysiswas only 45% of that of young leaves; (e) The expression of *PSK* after enzymolysis was only 2% of that of young leaves; (f) The expression of *CESA* after enzymolysis was only 47% of that of young leaves; (g) The expression of *GAUT* after enzymolysis was only 65% of that of young leaves.

## 3. Discussion

### 3.1 Effect of enzymatic digestion on the subcellular structure of sugarcane protoplasts

The level of viability is usually used as a criterion to determine the quality of the protoplasts after enzymatic digestion. However, enzymatic digestion generates structural variability resulting in anomalies observed during cell division as well as in plants regenerated from protoplasts (Cambecedes et al., 1988). These anomalies often hinder the introduction of new plant varieties obtained by methods of in vitro protoplast fusion (Handley et al., 1986). Our team found that the highly viable sugarcane protoplasts obtained by enzymatic digestion of young sugarcane leaves using the optimal mannitol concentration showed severe browning at a later stage and the cells were unable to divide continuously, which greatly hindered the regeneration of sugarcane protoplasts (Li. Z. G et al., 2020). Although, high yield (5 × 10^6^ protoplasts/g FW) and high vitality protoplasts (above 90%) were obtained by optimizing enzymatic hydrolysis conditions, The cell membranes of protoplasts perforate to different degrees after enzymolysis. Investigation of the tubulin cytoskeleton of protoplasts isolated from Medicago sativa and Nicotiana tabacum indicates that the perinuclear and radial cytoskeleton significantly limit the ability to proper cell division. These play a key role in the migration of the nucleus to the center of the cell and the maintenance of the proper position ofthe nucleus just before division (Meijer & Simmonds, 1988). In this research, the nucleolus of nuclear membrane was still intact after enzymatic hydrolysis, but the blue fluorescence and nuclear activity were weakened. Before enzymolysis, microtubules were tightly connected to the plasma membrane in young sugarcane cells, and a large number of periplasmic microtubules stuck to the plasma membrane of all newly isolated protoplasts in a fan-like pattern. The results indicate enzymatic digestion effect on the subcellular structure of sugarcane protoplasts and may be the cytological reason why highly viable protoplasts were also difficult to regenerate plants from highly viable sugarcane protoplasts.

### 3.2 Osmotic stress and oxidative stress occurred during enzymolysis

The successful isolation of protoplasts depends on breaking down the cell wall and releasing the intact protoplasts (J. Li et al., 2018). The enzymatic process requires the addition of mannitol and sucrose to regulate cellular osmolality. Higheror lower osmotic pressure can cause osmotic stress in protoplasts and reduces protoplast viability (Bai et al., 2020). The majority of the protoplast may exist but will not fracture (J. Chen et al., 2015). Even the protoplasts isolated from the solution were almost disintegrated (He et al., n.d, 1996). The possible reason is that during enzymolysis, because of the unstable cellular osmotic environment, while the cytoplasm is fragile and unevenly concentrated, the protoplasm breaks easily, affecting its ability to divide causes difficulty in regeneration of protoplasts (Z. Wu et al., 2021).

Osmotic stress causes multiple effects not only on cell physiology, but also on protein and gene expression. The osmotic stress effect at the time of protoplast separation altered the expression of some of the resistance genes leading to browning. For example, decreased expression of *DREB* resistance genes and repression of the *DREB / CBF-COR* pathway reduces plant tolerance to various abiotic stresses (Hayat et al., 2022). The cell wall is a very important defense system of plant cells, and enzymatic removal of the cell wall not only produces osmotic stress on the protoplasts (D. Wang et al., 2022), but also inevitably alters the expression of *NAC* secondary wall thickening promoter (NST)/secondary wall-associated *NAC* structural domain protein (*SND*) and *SOMBRERO (SMB*) subfamily proteins, thus making it difficult for *NST, SND*, and *SMB* to participate in SCW (secondary cell wall) forming cells (Kim et al., 2021). When the protoplast activity was higher than 70%, the purity and integrity of extracted RNA were higher (Li. S. L et al., 2019). In our research, the expression levels of stress-resistant genes *DREB, WRKY, MAPK4* and *NAC* were significantly up-regulated. It is suggested that increased osmotic pressure reduces cell volume and results in an increased concentration of ions and macromolecules, and show that several multivalent proteins and genes remain dispersed at physiological conditions and reversibly condense to microscopic granules during enzymolysis (Majumder & Jain, 2020).

Osmotic stress generally leads to an increase in ROS and induces oxidative stress (Noctor et al., 2015). Our research shows that the content of MDA increased significantly, 4.7 times that of young leaves, and the content of antioxidant anion O^2-^ decreased significantly, only 1.2% of that of young leaves. Reduces the expression of the resistance gene *MAPK*, which in turn reduces and inhibits the content and activity of the antioxidant enzyme system enzymes CAT, POD and SOD reduced viability and death (Apel & Hirt, 2004).

Our research shows that the activities of antioxidant enzymes POD, CAT and APX decreased to 17.7%, 6.5% and 17.5% respectively. The expression levels of *Gu/ZnSOD* and *CAT* were lower than those of young leaves, only 1.6% and 2.8% respectively. Sugarcane heterozygous cells were subjected to oxidative stress with significant changes in ROS and a massive accumulation of MDA in protoplasts. However, after enzymolysis, the protoplasmic ROS content was dramatically reduced in contrast to the previous ones, probably because the SOD activity was maintained at a high level and catalyzed a large amount of O^2-^. Studies have shown that ROS is one of the causes of protoplast browning, reduced viability and death (Apel & Hirt, 2004). SOD can disproportionate superoxide anions into hydrogen peroxide and oxygen, and increasing the activities of antioxidant enzymes such as SOD, CAT, POD and APX can reduce oxidative stress in maize (Zea mays) (Ma et al. 2015), reduce membrane damage during enzymolysis of peanut (Arachis hypogaea) protoplasts (He et al., 1994), and scavenge free radicals in tomato (Solanum lycopersicum) (Bai et al., 2020). During the process of protoplast isolation, cell wall removal and the abiotic stress it generates leads to a great decrease in the content and activity of antioxidant enzymes, which results in the inability to achieve a dynamic balance between intracellular ROS production and scavenging (S. Chen et al., 2006), which in turn leads to protoplast browning.

### 3.3 Effect of enzymatic digestion on the proteomics of sugarcane protoplasts

Abiotic stresses generated by enzymatic processes in sugarcane young leaves can cause continuous trauma to protoplasts. Because plants are constantly threatened by wounding throughout their lives, understanding the biological responses to wounds at the cellular level is critical (Son et al., 2021). Plant protoplasts constitute unique single-cell systems that can be subjected to genomic, proteomic, and metabolomic analysis (X. Xu et al., 2021). There are also proteomic studies showing that proteins such as ascorbate peroxidase (Holmes et al., 2006), dehydroascorbate reductase, glutathione transferase and mitochondrial manganese superoxide dismutase (Shi et al., 2008) are also associated with higher cytokinesis activity. Xiaoqin Wang (2017) has examined global changes in the proteome following protoplast development, using *iTRAQ* proteomic strategies coupled with *LC-MS/MS* (Y. Wang et al., 2017). One hundred andsixty-two proteins involved in defence responses, energy production, translation, metabolism, protein destination and storage, transport, transcription, cell growth/division, cell structure and signal transduction were identified. The study reveals the integration of the developmental reprogramming response in P. patens. Jinping Zhao (2019) used a label-free quantitative proteomics approach to analyze protein accumulation profiles of protoplast and chloroplast from RSV-infectedy established a method to elucidate the localization change of nucleus-encoded *ChRPs* and indicate a new layer of RSV–host interaction where the targeting of nucleus-encoded *ChRPs* is hindered during RSV infection. Our proteomics results showed that a total of 2,287 DEPs were identified after enzymatic digestion of young sugarcane leaves. Of which, 810 were up-regulated and 1,477 were down-regulated. The main biological processes of the major differentially expressed proteins included cellular processes (1,101 proteins), metabolic processes (1,049 proteins), stimulus responses (320 proteins), bioregulation (267 proteins), and cellular component organization or biogenesis (259 proteins), showing that these proteins are involved in dynamic networks in response to enzymatic digestion.

### 3.4 The expression of oxidation genes and protoplast regeneration genes was affected by enzymatic hydrolysis

Oxidative stress causes effects on gene expression in addition to cellular physiological and biochemical metabolism. Knocking out or reducing the expression of *SlMAPK3* will inhibit the activities of antioxidant enzymes (APX, POD, SOD and CAT) and induce the accumulation of H_2_O_2_ (Shu et al., 2022). However, osmotic stress caused by enzymatic hydrolysis of young sugarcane leaves makes the production and elimination of ROS in cells unable to maintain a stable state, resulting in the accumulation of ROS, which is the main reason for the browning and death of protoplasts (Khatri & Rathore, 2022). Research shows that, peroxisome proliferation and CAT contribute to ROS homeostasis and subsequent protoplast division induction (Tiew et al., 2015); Overexpression of *TaWRKY46* in wheat resulted increasesuperoxide dismutase (SOD), as well as higher activities of catalase (CAT) and peroxidase (POD) for regulating the osmotic balance and ROS scavenging (Yu & Zhang, 2021). Our research show that sugarcane heterozygous cells were subjected to oxidative stress during enzymatic digestion and most of the oxidase-related proteins and genes expression were down-regulated. *Gu/ZnSOD* and *CAT* expression were significantly down - regulated (Fig. 5), as well as most of DEPs such as ascorbate peroxidase, glutathione peroxidase, peroxidase, and catalase. The results showed that protoplasts were not able to handle oxidative stress well after enzymolysis.

Enzymatic hydrolysis of young sugarcane leaves reduces the expression of protoplast regeneration genes, which is also a major obstacle to protoplast regeneration. *CyclinD3*, *cyclinA*, *CyclinB*, and *CyclinE* regulated cell cycling for the cell proliferation (Ahn et al., 2018); Transcriptional levels of *CycD2* and *CDC2* (two genes regulating the progress of cell cycle) decreased, which inhibited the activity of vacuole invertase after the recovery of heat stress, thus shortening the cell length (Luo et al., 2021); *Phytosulokine, (PSK*) is a plant hormone involved in the information transmission between plant cells, and its expression decline will inevitably affect the development and growth of plants (de Souza et al., 2021); Seed plants use different *CESA* isoforms for primary and secondary cell wall deposition (X. Li et al., 2022); The *GAUT* gene family may affect cotton fiber development, including fiber elongation and fiber thickening (Senmiao et al., 2021). Our research shows that the expression levels of *CyclinD3, CyclinA, CyclinB, cdc2, PSK, CESA, GAUT* genes related to plant regeneration are significantly down-regulated after enzymatic hydrolysis of young sugarcane leaves, which are 52%, 21.32%, 18.60%, 45%, 2%, 47%, 65% of those of young sugarcane leaves, respectively. Compared with young leaves, these key genes have changed significantly after enzyme digestion. It is thought that the way enzymes break down somatic cells may have some effect on the way hybrid cells regenerate.

Technological breakthroughs in single-cell transcriptomic approaches have aided investigations of expression dynamics and biological processes at the cellular level (Mo & Jiao, 2022). In the study of plant molecular biology processes, the establishment of a suitable protocol for protoplast transformation will not only allow a detailed analysis of early signs of protoplast regeneration (e.g. chloroplast division and cell wall reconstruction), but also expand the prospects for functional studies of plants (Neubauer et al., 2022). Fluorescent dye labeling and the use of qPCR techniques can also be applied to protoplasts (Pasternak et al., 2005) for the analysis of abiotic stress on the expression of protoplast-related genes after enzymatic digestion. Our study showed that enzymatic digestion does produce osmotic stress on the protoplasts, and the protoplasts respond to the stress accordingly. The expression of resistance and regeneration genes were significantly up-regulated or down-regulated. Then, in future work, we can also use the protoplast gene transient expression system to locate the location and function of differential proteins during enzymatic digestion, which may be able to resolve to some extent the mechanism of the difficulty of protoplast regeneration due to changes in differential proteins of protoplasts during enzymatic digestion (such as changes in antioxidant enzyme activity). In addition, enzymatic digestion affects osmotic, oxidative and regenerative genes, and it is also possible to establish molecular markers for enzymatic digestion of protoplasts, calibrate the degree of enzymatic digestion, and screen the conditions of enzymatic digestion (selection of materials, composition of enzymatic solution, osmotic pressure, time, concentration, etc.) with a view to obtaining protoplasts with high yield quality and laying a solid foundation for obtaining regenerative plants (Figure 9).

**Figure 9.**
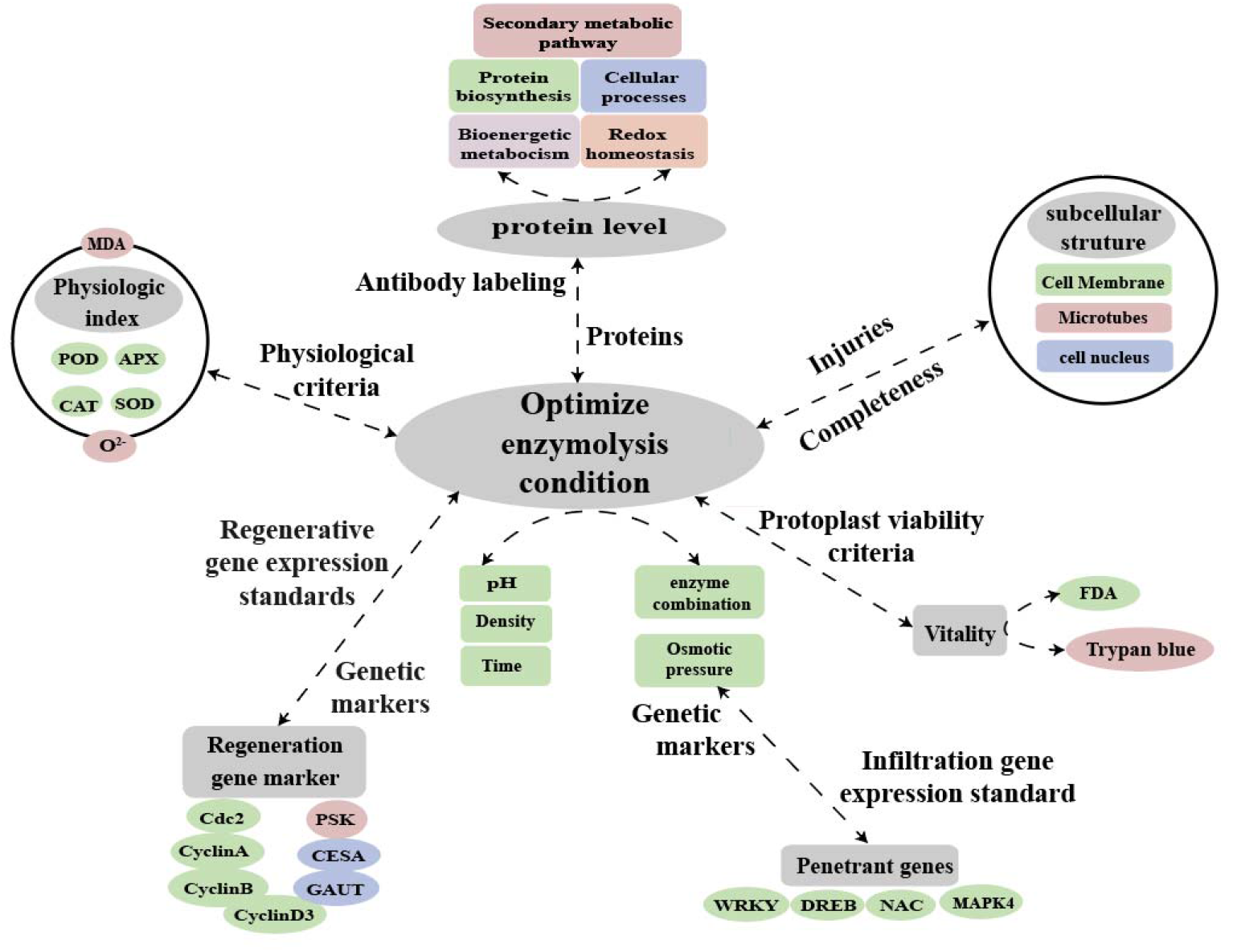
Hypothetical model for molecular labeling and detection of antibody of protoplast during enzymatic digestion.

## 4. Conclusion

Our research shows that enzymatic hydrolysis does produce osmotic stress and oxidative stress on protoplasts, which causes the sugarcane protoplast proteomics (bioenergetic metabolism, cell wall synthesis, cell cycle regulation, and oxidative stress after enzymatic digestio) differentially expressed, the resistance to stress related enzyme activity (SOD, POD, CAT, and APX) and remove oxidation product(s O^2-^) were declined, the physiological indexes of oxidation products(MDA) were improved, the expression of genes (*GAUT, CESA, CyclinA, CyclinB, CyclinD3, cdc2* and *PSK*) related to protoplast regeneration is different from that of sugarcane young leaves.

Altogether, this study provides a better understanding of the molecular, physiological and cytological mechanism about the difficulty in sugarcane protoplast regeneration Figure 10), and provides parameters to establish a standard system for obtaining regenerated protoplasts using molecular markers and antibody detection of enzymolysis in the future.

**Figure 10:**
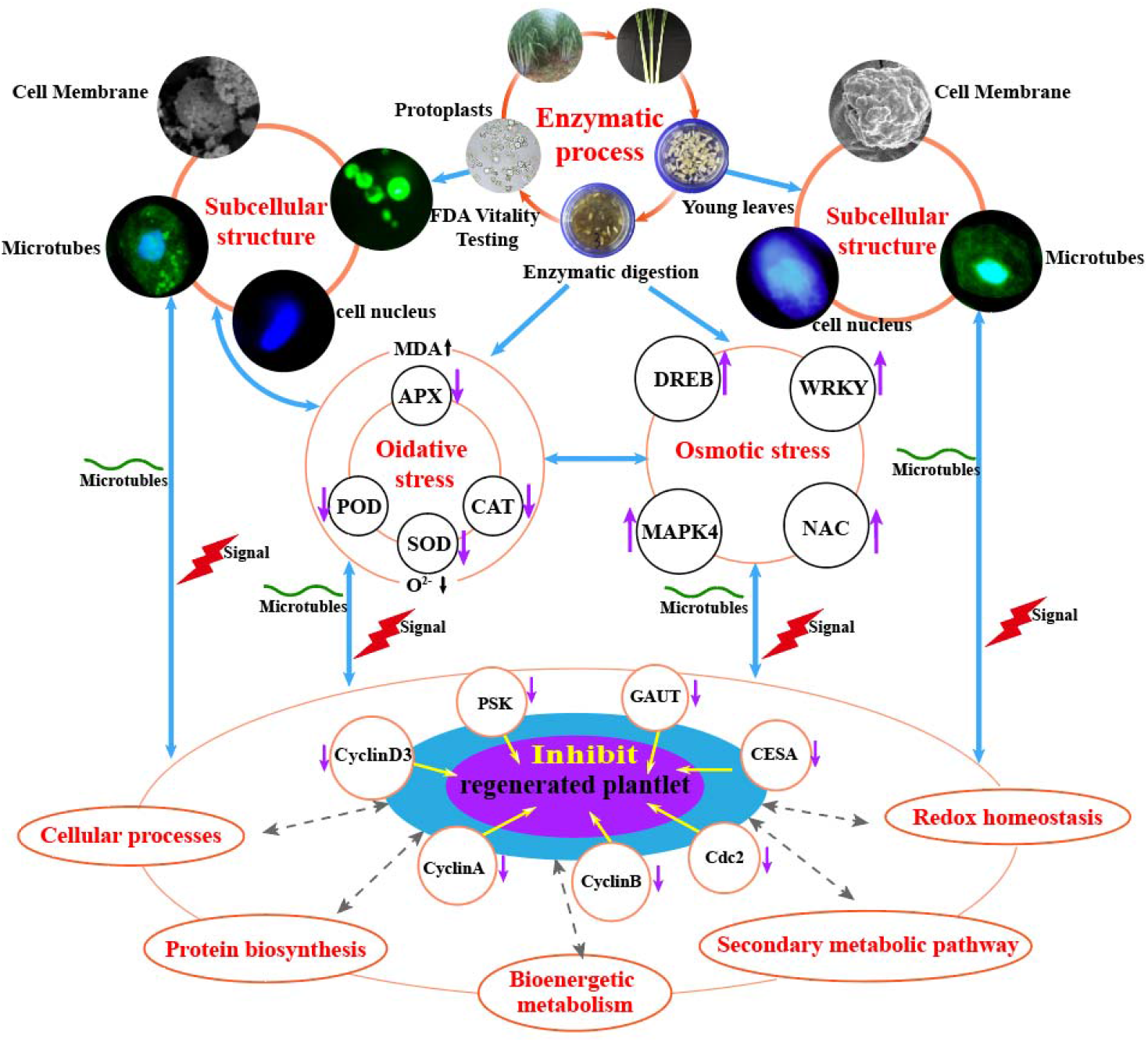
Molecular and physiological mechanisms of enzymatic hydrolysis affecting the difficulty of sugarcane protoplast regeneration.

## 5. Materials and Methods

### 5.1 Plant material, protoplast material acquisition

The sugarcane variety ROC22 planted in the experimental base of sugarcane (Saccharum offcinarum) in the College of Agriculture of Guangxi University was used as the material, and the young leaves were sampled at an early elongation stage. For further use, the samples were quickly frozen in liquid nitrogen and sorted at 80 °C.

The separation of protoplast from young sugarcane leaves was done based on the method proposed by Zhang (2022) (D. Zhang et al., 2022). The robust tail sheaths of sugarcane at the initial stage of elongation were selected as the enzymolysis materials of young leaves. The outer 2 - 3 layers of the leaf sheaths were peeled off first and then sterilized with 75% alcohol for 30 s. The outer layer and leaf sheaths at both ends were removed after three washes with sterile water to reveal the light-yellow central leaves. The young leaves 1 - 5 cm above the growing point were cut into slices with a thickness of about 1 mm. 0.5 g of young leaves were collected and put into 5 mL of CPW solution (containing 13% mannitol, pH 5.8). After plasmolysis for 0.5 - 1 h, the CPW (containing 13% mannitol) solution was removed and 5 mL of enzymolysiss solution was added to perform the enzymolysis at room temperature for 4 h. The protoplast suspension was then filtered through 100 and 200-mesh cell sieves, and the protoplasts were purified by gradient centrifugation.

### 5.2 Protoplast viability testing

The FDA stock solution was dissolved in acetone to 5 g/L and stored at −20 °C. The FDA working solution was made by diluting the above stock solution with PBS to 5 mg/L. Suspend 1 × 10^6^ protoplasts in 0.01 mL FDA working solution, stain at 37 °C for 5 min, add PBS to 0.1 mL.

The total number of cells in the samples measured in this study was 2.5 × 10^6^ g/FW. Protoplasmic vitality (%)= Number of stained cells / total number of cells×100%

### 5.3 Characterization by scanning electron microscopy

Protoplasmic SEM sample preparation and equipment parameters were referenced to Min Lu et al. (Lu et al., 2021)

### 5.4 Nucleus morphology of young sugarcane leaves and protoplasts

The chromosomes were examined in a dark room by applying 30 μl DAPI (0.5 μg/ml) drops per sheet to the microscopically examined chromosome films, covered with coverslips, observed by Leica DMRA2 fluorescence microscope, Leica QFI the images were captured by the SH DC 350F camera system, and the average pixel values of chromosomes were measured by Leica cw 4000.

### 5.5 Changes in the microtubule skeleton of sugarcane young leaves and protoplasts

Frozen section method combined with indirect immunofluorescence technique and DAPI staining was used to observe the arrangement of the cellular microtubule array of sugarcane young leaves and protoplasts using fluorescence microscopy (refer to Li Zhigang et al.) (Li. Z. G et al., 2008).

### 5.6 Sample collection and processing for proteomics analysis

Young leaves: The sugarcane ROC22 was used, and the young leaves were sampled at early elongation stage. The samples were quick-frozen in liquid nitrogen and sorted at −80 °C for further use. Three replicates were set. Protoplasts: After ROC22 enzymolysis, the density of protoplasts was adjusted to 1 × 10^6^/mL, 0.5 mL were taken and shaken well, then frozen in liquid nitrogen and stored at −80 °C. Three replicates were set.

### 5.7 Protein Extraction, Digesting, and iTRAQ Labeling

The experimental method was referenced to Wang et al. (2022). A 0.5 g sample of sugarcane young leaves and 5× 10^6^ protoplasts were homogenized in 2 mL of lysis buffer consisting of 8 M urea, 50 mM Tris-HCl (pH 8), 0.2% SDS and sonicated on ice for 5 min, respectively. The samples were then centrifuged at 12,000× *g* at 4 °C for 15 min, and the supernatant was moved to a new tube. The concentration of protein was determined using a Bradford protein assay. Extracts from each sample were reduced with 2 mM DTT for 1 h and alkylated with sufficient iodoacetic acetate for 1 h in the dark at room temperature. A 4-fold volume of precooled acetone was mixed with the samples and incubated at 20 °C for 1 h; the samples were then centrifuged to collect precipitation. After they were washed three times with cold acetone, the pellets were dissolved in lysis buffer containing 0.1 M triethyl ammonium bicarbonate (TEAB, pH 8.5) and 8 M urea. An equal number of proteins were digested with trypsin (Promega, Madison, WI, USA) at a ratio of 1:50 (w:w) for 16 h at 37 °C and iTRAQ reagents kit (Applied Biosystems, Framingham, MA, USA) were used to label 100 micrograms of digested proteins according to the manufacturer’s protocol.

### 5.8 HPLC Fractionation and LC-MS/MS Analysis

A SHIMADZU LC system equipped with a 20 AB column (Gemini C18 4.6 × 250 mm, 5 μm) was used for the high performance liquid chromatography analysis. LC-MS/MS testing was conducted by UltiMate 3,000 UHPLC (thermo, Massachusetts, USA). The protein quantification assay was done based on the method proposed by Xu Ke (K. Xu et al., 2022).

### 5.9 Real-Time Quantitative PCR Analysis of Proteins

Total RNA extraction and reverse transcription of RNA into cDNA were performed based on the method described as manufacturer’s protocol of Prime Script TMΠ 1stStrand cDNA Synthesis Kit andTB GreenTM Premix Ex TaqTM II (TaKaRa, Beijing, China). qRT-PCR was performed with ChamQ Universal SYBR qPCR Master Mix (Vazyme, Nanjing, China) and the QuantStudio 5 Real-time PCR system (Applied Biosystems, Maltham, MA, USA) (K. Xu et al., 2022).

### 5.10 Proteomic Analysis

Functional analysis of identified proteins was performed using Gene Ontology (GO)(http://www.geneontology.org). Differentially accumulated proteins were then placed in the KEGG database (http://www.kegg.jp/kegg/pathway.html). In order to determine functional subgroups and metabolic pathways in which the differentially accumulated proteins were enriched, GO, and KEGG pathway enrichment analyses were conducted. The cluster analysis of the differentially accumulated proteins was conducted using Cluster 3.0, and the heatmap was generated using TreeView version 1.6.

### 5.11 Functional analysis of DEGs

Functional analysis was performed on all genes with significant,differences in expression using the gene ontology (GO) and Kyoto Encyclopedia of Genes and Genomes (KEGG) databases, based on the principle of hypergeometric distribution. Omic Share was used to perform cluster analysis and GO and KEGG function enrichment analysis.

### 5.12 Determination of enzyme activity

In this experiment, the enzyme activity was measured by the conventional method. POD activity was determined by guaiacol method; APX and CAT using the UV absorption method. Superoxide dismutase is a family of metallo enzymes which is known to acceleraie spontaneous dismutation of the superoxide radical to hydrogen peroxide and molecular oxygen. SOD is widely distributed among aerobically living organisms and has been inferred to play an important role incontrolling superoxide levels in cellular compartments (Paoletti et al., 1986).We refer to the Francexo Paoletti method,which use a stable and inexpensive reagent for rapid and highly sensitive measurement of SOD activity in pure enzyme preparations.

## Supplemental data

Supplementary Table S1. Candidate DEPs associated with energy metabolism

Supplementary Table S2. Candidate DEPs associated with cell walls

Supplementary Table S3. Candidate DEPs associated with cell cycle

Supplementary Table S4. Candidate DEPs associated with synthesis of secondary metabolites

Supplementary Table S5.Candidate DEPs associated with antioxidant

## Acknowled gments

The authors thank the College of Agriculture, Guangxi University, Guangxi Key Laboratory of Sugarcane Biology for providing a good research platform and its support for experimental techniques.

